# Widespread Prevalence of CD19 Exon 5-6 Skipping In Indian Pediatric B-cell Acute Lymphoblastic Leukemia Patients

**DOI:** 10.1101/2024.10.15.618619

**Authors:** Devesh Srivastava, Anurag Gupta, Nishant Verma, Ashish Misra

## Abstract

B-cell acute lymphoblastic leukemia (B-ALL) is characterized by the malignant burgeoning of abnormal B-cell lymphoblasts. In recent years, the use of CART therapy which targets CD19 antigen present on the surface of B-cells, has gained significant attention as a treatment option against aggressive and refractory forms of B-ALL. However, the loss of CD19 antigen on B-cell surface due to aberrant splicing under therapy pressure has been suggested as one of the main factors for the emerging CART therapy resistance. Herein, using RT-PCR based splice assays we examined CD19 splicing patterns in 43 primary pediatric B-ALL patient samples spread across various subtypes. We observed that CD19 isoform lacking exon 5-6 exists in ∼ 55% of pediatric patients at the initial diagnosis stage itself. Using in-silico analysis, we identified RNA binding proteins, RC3H1 and MBNL1, as potential regulators of exon 5-6 skipping. Furthermore, qRT-PCR analysis in patient samples revealed that RC3H1 and MBNL1 are significantly upregulated in samples exhibiting exon 5-6 skipping. Taken together, we for the first time report the existence of aberrantly spliced *CD19* isoform lacking exon 5-6 in primary pediatric patients, and this occurrence could potentially result from RC3H1 and MBNL1 dysregulation.

## Introduction

B-cell acute lymphoblastic leukemia (B-ALL) is a harmful blood malignancy arising from the uncontrolled proliferation of immature B-cell lymphoblasts in the bone marrow, leading to fatal consequences in both pediatric and adult populations ^1,2^. Notably, B-ALL is the most common form of leukemia observed in children as around 80% of leukemia cases in children under the age of 15 are of B-ALL origin ^3^. In last few decades, using conventional combination chemotherapy, remission rate of more than 90% has been observed in B-ALL patients ^4,5^. However, due to heterogeneity amongst different B-all subtypes, chemo resistance, aggressive relapse and eventual death is also observed in many patients. The use of chimeric antigen receptor (CAR)-T cells (CART) therapy, a form of immunotherapy, as a treatment option against aggressive and refractory forms of B-ALL, has gained momentum in recent years ^6,7^. In CART therapy for B-ALL, T-cells are genetically modified in a laboratory to express a CAR that specifically targets the CD19 antigen found on the surface of all leukemic B-cells. Subsequently, these modified T-cells are infused back into the patient’s body intravenously, to effectively target and eliminate leukemic cells ^8^ (rewrite).

Human CD19 gene contains 15 exons and is located on chromosome 16p11.2. It encodes for a 95kda transmembrane glycoprotein ^9^ which contains extracellular Ig domains, a transmembrane domain, and an long cytoplasmic domain ^9,10^. CD19 is a pan B-cell antigen that is expressed in the earliest stages of B-cell development and is necessary for normal differentiation and maturation of B-cells ^11^. Moreover, CD19 is a crucial marker for lineage assignment of blasts as per World Health Organization’s (WHO) guidelines (WHO blue book) and is present in almost all cases of B-ALL, making it the perfect target for CARTs. Initially, CART treatment demonstrated remarkable and encouraging outcomes, achieving a remission rate of approximately 90% in B-ALL patients ^10^. However, over time, post CART therapy, disease relapse was observed in ∼ 30-50% of the patients ^12,13^. In most cases, relapse is attributed to the loss of the CD19 surface antigen, rendering the malignant B-cells unrecognizable to CARTs ^12,14^. Various splicing events, such as deletion, skipping, and retention of specific exons & introns, have been suggested as processes causing damage to the CD19 epitope with many of these events being present prior to any therapeutic interventions. Sotillo et al., studied relapsed CART patient samples and proposed that skipping of CD19 exon 2 and exon 5-6 can lead to loss of CD19 surface antigen, contributing towards CART resistance ^15^. Moreover, Fischer et al., analyzed primary B-ALL patient samples and reported that CD19 skipped exon 2 isoform can exist at diagnosis stage itself ^16^. Likewise, CD19 intron 2 retention has also been reported in CART relapsed and primary B-ALL patient samples ^17,18^. Currently, literature lacks studies on the existence of alternatively spliced isoforms with exon 5 and 6 exclusions in de novo primary pediatric B-ALL. Exons 5 and 6 of the CD19 protein encode for its transmembrane & cytosolic domains, and their loss is predicted to result in an altered CD19 isoform which can escape detection by CARTs (Figure 1a) ^16,19^.

**Figure 1.**
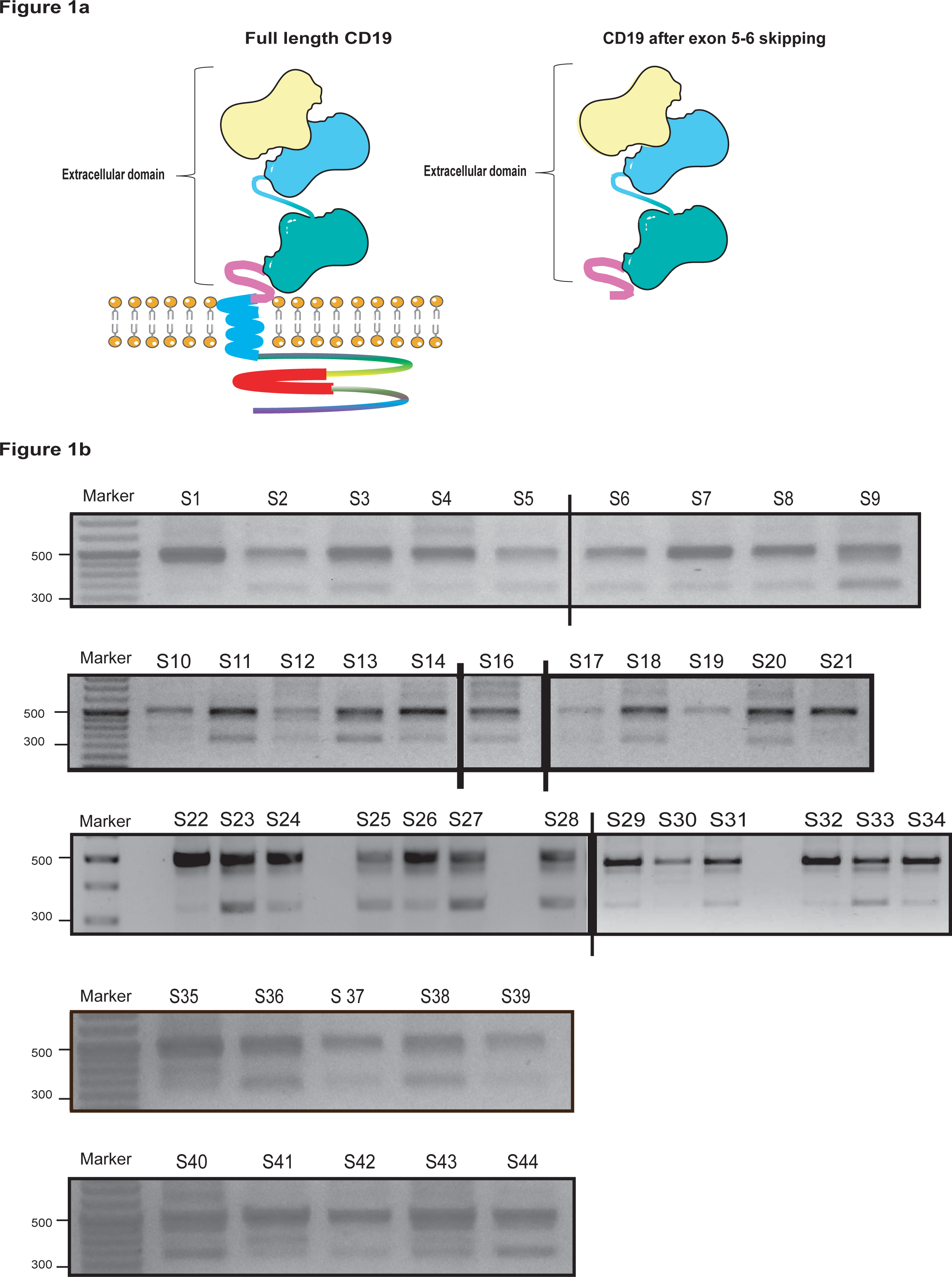
Full-length CD19 and truncated CD19 isoform. (a) The domain structure of the full-length CD19 protein includes extracellular, transmembrane, and cytosolic domains. Exons 5 and 6 are responsible for encoding the transmembrane and cytosolic domains, respectively. Skipping these exons leads to the production of a truncated CD19 protein. (b) CD19 skipped exon 5-6 (SE5-6) isoform detection using semi-quantitative RT-PCR in primary pediatric B-ALL patients. Here 490bp band represents full-length CD19 whereas 330bp band represents CD19 SE5-6 isoform. Out of 43 patient samples of different subtypes, 25 samples had intense or faint SE5-6 bands along with full-length bands while 18 samples had only full length and no SE5-6 bands.

In this study we for the first time have analyzed the skipping of CD19 exon 5 and 6 (SE5-6) isoform in 43 cases of de novo pediatric B-ALL. Furthermore, using in-silico analysis we have identified the RNA-binding proteins (RBPs) that bind to the exon 5-6 loci and can potentially influence the skipping of exon 5-6. These RBPs are significantly upregulated in patients exhibiting exon 5-6 skipping.

## Materials and Methods

### Patient sample collection and processing, FISH data analysis

This study comprised of 43 cases of de novo pediatric B-ALL patients. Ethical approvals were obtained (IEC Protocol No. IITH/IEC/2022/07/12) from the participating institutes. Bone marrow aspirate and peripheral blood samples were collected from patients with informed consent. Peripheral blood counts were recorded, and morphological assessments of blood & bone marrow aspirates were conducted to determine blast percentages. Patients with blast counts < 30% were excluded from this study. Samples underwent flow cytometric immunophenotyping using a BD FACSLyric instrument with antibodies targeting B, T, and myeloid lineage markers. The study included patients aged 1-12 years, while those under 1 year were excluded due to sample size constraints. Fluorescence in situ hybridization (FISH) was used to detect genetic abnormalities including BCR::ABL1, ETV6::RUNX1 fusions, KMT2A rearrangements, TCF3, ZNF384, CRLF2, CSF1R, PDGFRB, JAK2, ABL1 and ABL2. Samples negative for WHO defined abnormalities were classified as B-others.

### RNA isolation, cDNA preparation and PCR

RNA was extracted from the peripheral blood and/or bone marrow samples using QIAamp RNA Blood Mini Kit (Qiagen, Hilden, Germany) as per the manufacturer’s instructions. Following RNA isolation, cDNA preparation was done with 2ug of RNA using ProtoScript II Reverse Transcriptase (New England Biolabs) as per manufacturer’s instructions. To identify the SE5-6 transcripts in CD19, the following primer pair spanning exon 4 to exon 8 was used, forward: 5’ AAGGGGCCTAAGTCATTGCT 3’ and reverse: 5’ TGCTCGGGTTTCCATAAGAC 3’, as described by Sotillo et al ^15^. CD19 PCR of cDNA samples was done using 12.5ul of Amplitaq gold 360 mastermix (Thermo Scientific, Dreieich, Germany), 2uL of cDNA, 0.5uL of each forward and reverse primers and 4.5uL of nuclease free water per reaction under following cycling conditions: Initial denaturation at 95°C for 10 mins, followed by 35 cycles of denaturation 95°C for 30 secs, annealing 62°C for 30 secs, extension 72°C for 30 secs, and final extension of 72°C for 7 mints.

### Sequencing

Sanger sequencing was performed to confirm the nature of the bands obtained on gel electrophoresis. The bands were cut under UV transillumination. The band was later eluted from the gel using NucleoSpin Gel and PCR cleanup kit (Macherey-Nagel). The eluted product was cleaned using ExoSAP-IT™ (Thermo Scientific, Dreieich, Germany) and sequenced in Applied Biosystems 3500 genetic analyzer using BigDye™ Terminator v3.1 Cycle Sequencing Kit (Thermo Scientific, Dreieich, Germany). Germany). The data was analyzed and aligned with the CD19 transcript using Geneious Primer version 2023.2.1 software.

### In-silico analysis

To identify the RBPs interacting with CD19 exon 5-6 (chr16:28935221-28936411) locus we performed CD19 sequence analysis using RBPmap (https://rbpmap.technion.ac.il/) and oRNAment (https://rnabiology.ircm.qc.ca/oRNAment). RBPmap uses a weighted-rank technique to map the motifs present on RNA sequences and allows the user to choose their desired motifs ^20^. oRNAment uses a search algorithm which provides a matrix similarity score between 0 an1 after scanning each transcript ^21^. To shortlist RBPs, we had set the stringency level to highly stringent in the RBPmap and for oRNAment we selected RBPs with score > 0.1.

### Patient data analysis

RNA-Seq RPKM file containing gene expression data of 203 B-ALL patients was retrieved from the project: “Pediatric Acute Lymphoid Leukemia - Phase II (TARGET, 2018), deposited in online cancer repository cBioPortal (https://www.cbioportal.org/) ^22^. Additionally, mRNA expression data for the selected RBPs was also retrieved from cBioPortal using the same above-mentioned project (mention study id or something).

### Statistical tests

Statistical tests (unpaired t-test) for qRT-PCR and expression analysis of shortlisted RBPs was done using GraphPad prism software. Correlation analysis of the shortlisted RBPs was also conducted using GraphPad Prism.

## Results

### Patient characteristics

Patient samples were collected at the time of initial diagnostic workup and all the patients included in the study exhibited CD19 expression. CD19 was moderately expressed in ∼70% cases, around ∼25% cases had dim to moderate CD19 expression, and less than 5% cases had low CD19 expression. The average age of the study population was ∼5 years. Of total, 62% were males while 38% were females. Hemoglobin of the study population varied from 3 gm/dL to 13 gm/dL with 96% of the patients presenting with anemia (Hb <11gm/dL) at diagnosis. Total leucocyte counts varied from 1 × 103 cells/ uL to 232 × 103 cells/ uL. 30% of the patients had leucopenia while 38% had leucocytosis. Except for one case, S37, all patients were thrombocytopenic. Blast percentages were higher in bone marrow samples as compared to peripheral blood. More than 50% blasts were present in peripheral blood and bone marrow in 80% and 93.5% of the cases respectively. Patient characteristics are summarized in table 1.

**Table 1.**
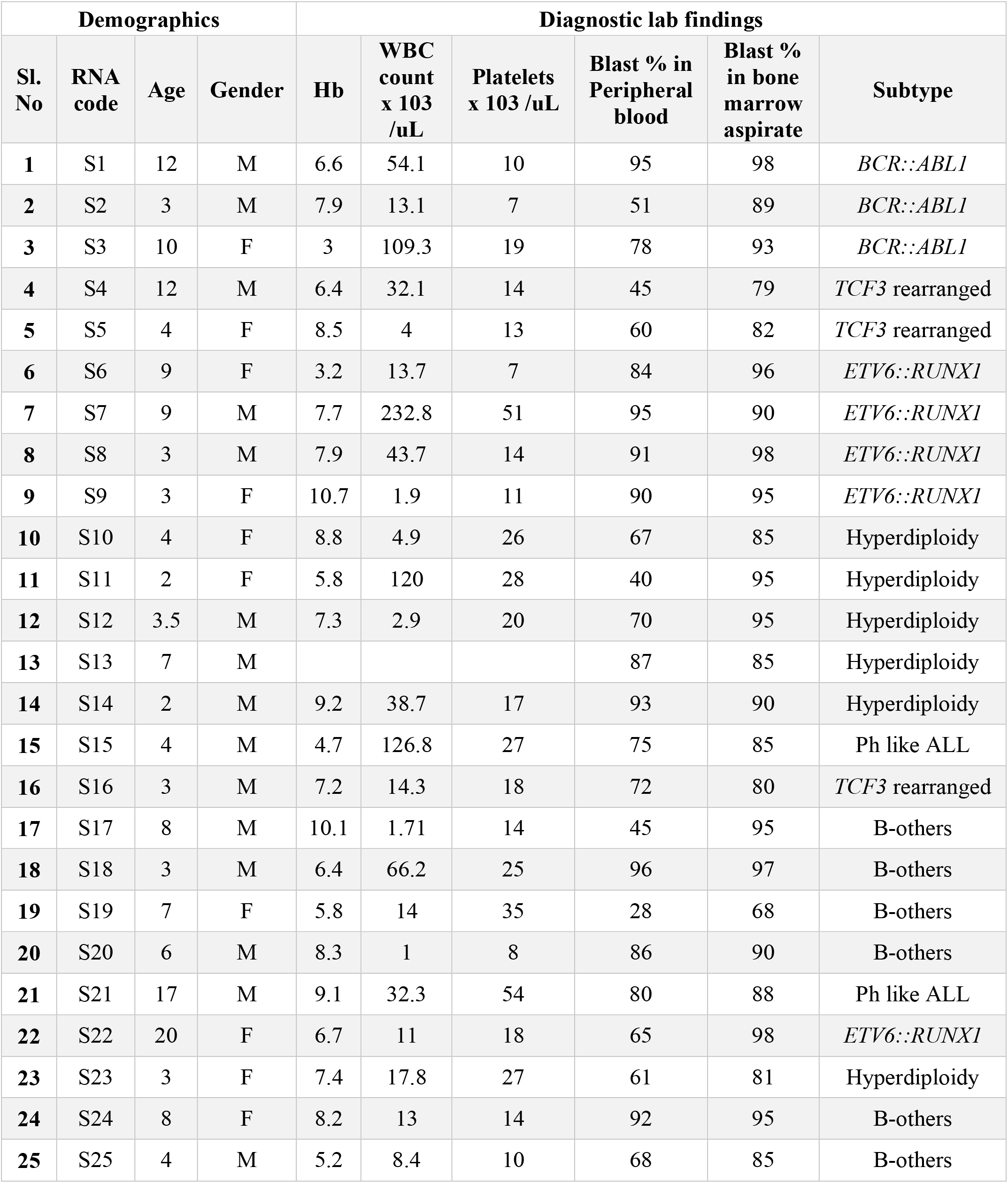
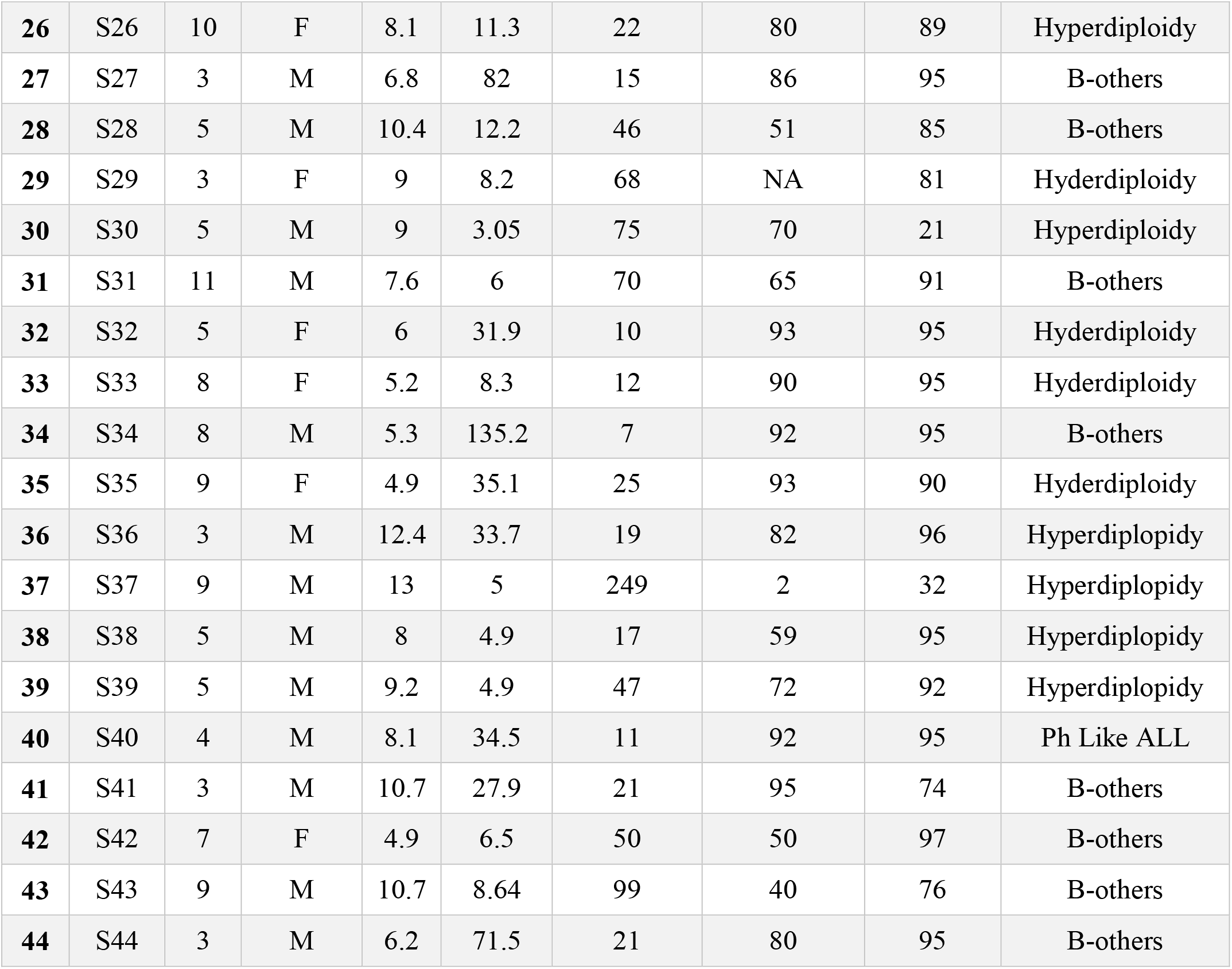
Diagnostic lab findings and patient characteristics.

| Demographics |  |  |  | Diagnostic lab findings |  |  |  |  |  |
| --- | --- | --- | --- | --- | --- | --- | --- | --- | --- |
| Sl. No | RNA code | Age | Gender | Hb | WBC count x 103 /uL | Platelets x 103 /uL | Blast % in Peripheral blood | Blast % in bone marrow aspirate | Subtype |
| 1 | S1 | 12 | M | 6.6 | 54.1 | 10 | 95 | 98 | *BCR::ABL1* |
| 2 | S2 | 3 | M | 7.9 | 13.1 | 7 | 51 | 89 | *BCR::ABL1* |
| 3 | S3 | 10 | F | 3 | 109.3 | 19 | 78 | 93 | *BCR::ABL1* |
| 4 | S4 | 12 | M | 6.4 | 32.1 | 14 | 45 | 79 | *TCF3* rearranged |
| 5 | S5 | 4 | F | 8.5 | 4 | 13 | 60 | 82 | *TCF3* rearranged |
| 6 | S6 | 9 | F | 3.2 | 13.7 | 7 | 84 | 96 | *ETV6::RUNX1* |
| 7 | S7 | 9 | M | 7.7 | 232.8 | 51 | 95 | 90 | *ETV6::RUNX1* |
| 8 | S8 | 3 | M | 7.9 | 43.7 | 14 | 91 | 98 | *ETV6::RUNX1* |
| 9 | S9 | 3 | F | 10.7 | 1.9 | 11 | 90 | 95 | *ETV6::RUNX1* |
| 10 | S10 | 4 | F | 8.8 | 4.9 | 26 | 67 | 85 | Hyperdiploidy |
| 11 | S11 | 2 | F | 5.8 | 120 | 28 | 40 | 95 | Hyperdiploidy |
| 12 | S12 | 3.5 | M | 7.3 | 2.9 | 20 | 70 | 95 | Hyperdiploidy |
| 13 | S13 | 7 | M |  |  |  | 87 | 85 | Hyperdiploidy |
| 14 | S14 | 2 | M | 9.2 | 38.7 | 17 | 93 | 90 | Hyperdiploidy |
| 15 | S15 | 4 | M | 4.7 | 126.8 | 27 | 75 | 85 | Ph like ALL |
| 16 | S16 | 3 | M | 7.2 | 14.3 | 18 | 72 | 80 | *TCF3* rearranged |
| 17 | S17 | 8 | M | 10.1 | 1.71 | 14 | 45 | 95 | B-others |
| 18 | S18 | 3 | M | 6.4 | 66.2 | 25 | 96 | 97 | B-others |
| 19 | S19 | 7 | F | 5.8 | 14 | 35 | 28 | 68 | B-others |
| 20 | S20 | 6 | M | 8.3 | 1 | 8 | 86 | 90 | B-others |
| 21 | S21 | 17 | M | 9.1 | 32.3 | 54 | 80 | 88 | Ph like ALL |
| 22 | S22 | 20 | F | 6.7 | 11 | 18 | 65 | 98 | *ETV6::RUNX1* |
| 23 | S23 | 3 | F | 7.4 | 17.8 | 27 | 61 | 81 | Hyperdiploidy |
| 24 | S24 | 8 | F | 8.2 | 13 | 14 | 92 | 95 | B-others |
| 25 | S25 | 4 | M | 5.2 | 8.4 | 10 | 68 | 85 | B-others |
| 26 | S26 | 10 | F | 8.1 | 11.3 | 22 | 80 | 89 | Hyderdiploidy |
| 27 | S27 | 3 | M | 6.8 | 82 | 15 | 86 | 95 | B-others |
| 28 | S28 | 5 | M | 10.4 | 12.2 | 46 | 51 | 85 | B-others |
| 29 | S29 | 3 | F | 9 | 8.2 | 68 | NA | 81 | Hyderdiploidy |
| 30 | S30 | 5 | M | 9 | 3.05 | 75 | 70 | 21 | Hyperdiploidy |
| 31 | S31 | 11 | M | 7.6 | 6 | 70 | 65 | 91 | B-others |
| 32 | S32 | 5 | F | 6 | 31.9 | 10 | 93 | 95 | Hyderdiploidy |
| 33 | S33 | 8 | F | 5.2 | 8.3 | 12 | 90 | 95 | Hyderdiploidy |
| 34 | S34 | 8 | M | 5.3 | 135.2 | 7 | 92 | 95 | B-others |
| 35 | S35 | 9 | F | 4.9 | 35.1 | 25 | 93 | 90 | Hyderdiploidy |
| 36 | S36 | 3 | M | 12.4 | 33.7 | 19 | 82 | 96 | Hyperdiploidy |
| 37 | S37 | 9 | M | 13 | 5 | 249 | 2 | 32 | Hyperdiploidy |
| 38 | S38 | 5 | M | 8 | 4.9 | 17 | 59 | 95 | Hyperdiploidy |
| 39 | S39 | 5 | M | 9.2 | 4.9 | 47 | 72 | 92 | Hyperdiploidy |
| 40 | S40 | 4 | M | 8.1 | 34.5 | 11 | 92 | 95 | Ph Like ALL |
| 41 | S41 | 3 | M | 10.7 | 27.9 | 21 | 95 | 74 | B-others |
| 42 | S42 | 7 | F | 4.9 | 6.5 | 50 | 50 | 97 | B-others |
| 43 | S43 | 9 | M | 10.7 | 8.64 | 99 | 40 | 76 | B-others |
| 44 | S44 | 3 | M | 6.2 | 71.5 | 21 | 80 | 95 | B-others |

### CD19 splicing gels exhibit alternatively spliced isoform

We performed semiquantitative RT-PCR in the primary pediatric B-ALL patient samples spread across different subtypes, using primers spanning exons 4-8. Using these primers, the expected band size for full-length wild-type CD19 transcript is 490 bp size whereas a transcript with SE5-6 is expected to give a band size of 330bp (exons 5 and 6 are 111bp and 49bp long, respectively). Strikingly, in our RT-PCR analysis, we observed that a significant percentage of patient samples demonstrated the presence of CD19 SE5-6 variant at the diagnosis stage itself, prior to any therapy (Figure 1b). The samples exhibited variation in the intensity of full-length & SE5-6 transcripts, with most samples showing prominent full-length transcript compared to SE5-6 transcript (Figure 1b). Next, we performed densitometric analysis to quantify CD19 full length and SE5-6 bands using ImageJ software. Based on densitometric analysis we set a threshold of 10% for SE5-6/full-length band intensity, above which a sample was considered to exhibit SE5-6 band while those below the threshold were considered as non-skipping samples (Supplementary figure 1a). We observed that out of our 43 patient samples, 25 samples exhibited the presence of SE5-6 band to varying degrees along with the full-length CD19 band, whereas 18 samples exhibited only full-length band and had no/background SE5-6 band. Furthermore, intense SE5-6 bands (SE5-6/full-length > 30%) were observed in around 25% samples such as: S9, S13, S23, S25, S27, S28, S33, S36, S38, S40 and S44. Notably, we observed that SE5-6 bands were present in primary patient samples irrespective of the subtype, i.e., the skipping was not subtype specific and also did not appear due to any therapy pressure.

To confirm the nature of the obtained bands, the full-length & SE5-6 bands were gel eluted and sequenced using the Applied Biosystems 3500 genetic analyzer. The data was analyzed and aligned with CD19 transcript using Geneious Primer version 2023.2.1 software. The sequencing result validated the presence of SE5-6 isoform. The 490 bp band representing the full-length transcript mapped precisely on to the full length CD19 transcript with no gaps. The 330 bp size band representing the SE5-6 transcript mapped to exons 4, exon 7 and exon 8 of CD19 transcript with a gap of 160 bp (exon 5 is of 111bp and exon 6 of 49 bp length – both skipped), thereby confirming the nature of the bands (Supplementary figure 1b).

### In-silico identification of RNA binding proteins and qRT-PCR analysis

RBPs are known to regulate exon inclusion and skipping by binding to exon splicing enhancer (ESE) sites in the pre-mRNA. Previous studies have reported that RBPs and related splice factors, regulate aberrant splicing events in CD19, for instance, Sotillo et al., reported that SRSF3 controls exon 2 inclusion in CD19 ^15^. Likewise, Ziegler et al., observed that PTBP1 modulates retention of intron 2 in CD19 ^18^. In addition, Cortés-López et al., also reported that PTBP1 regulates intron 2 retention in CD19 ^23^.. To identify the RBPs that might interact with CD19 exon 5-6 locus and affect splicing, we performed RBP analysis by exploring publicly available RBPmap ^24^ and oRNAment ^21^ databases. To select putative RBPs, stringency level was set to highly stringent in RBPmap, and oRNAment score > 0.1 was considered. We shortlisted the common RBPs obtained from these databases using VENNY 2.1 (https://bioinfogp.cnb.csic.es/tools/venny/). BOLL, LIN28A, MBNL1, NUPL2, PABPN1L, PCBP2, RC3H1, SNRPA, SRSF1, SRSF2, SRSF5, SRSF9 were the 12 common RBPs identified using VENNY 2.1 (Supplementary figure 2). RBPs with more than 5 binding sites in the exon 5-6 locus (from RBPmap-UCSC genome browser’s data) and significantly linked with B-ALL pathophysiology were further shortlisted. MBNL1, PCBP2, RC3H1 and SRSF2 each with more than 5 binding sites were selected for further analysis due to their increased likelihood of binding (Figure 2a). Further, to ascertain the correlation between expression levels of these four RBPs and B-ALL pathophysiology, we investigated “Pediatric Acute Lymphoid Leukemia - Phase II (TARGET, 2018)” data deposited in cBioPortal (https://www.cbioportal.org/). Our analysis revealed that expression levels of MBNL1, PCBP2, RC3H1, and SRSF2 is significantly elevated in B-ALL patients exhibiting any alteration (such as mutations, truncations, copy number variations) in these RBPs compared to patients with no alterations (Supplementary figure 3a-3d). Subsequently, using qRT-PCR, we analyzed the expression of these four RBPs across our pediatric patient samples (non-skipping vs SE5-6 samples) and observed a strong correlation between the expression levels of RC3H1 and MBNL1 with exon skipping. RC3H1 and MBNL1 were significantly upregulated in SE5-6 samples compared to non-skipping samples whereas PCBP2 and SRSF2 had no such change in their expression (Figure 2b-2c, Supplementary figure 4a-4b). Further, we extracted and analyzed RNA-seq RPKM gene expression file of 203 B-ALL patients using the same TARGET-phase II pediatric ALL dataset and found that RC3H1 and MBNl1 are significantly correlated with each other (Figure 2d). This correlation of RC3H1 and MBNl1 is in line with our findings which shows upregulation of both the RBPs in skipping samples. Our results imply that RC3H1 and MBNL1 might be potential regulators of exon 5-6 skipping in B-ALL patients, however, more in-depth analysis is required to substantiate it.

**Figure 2.**
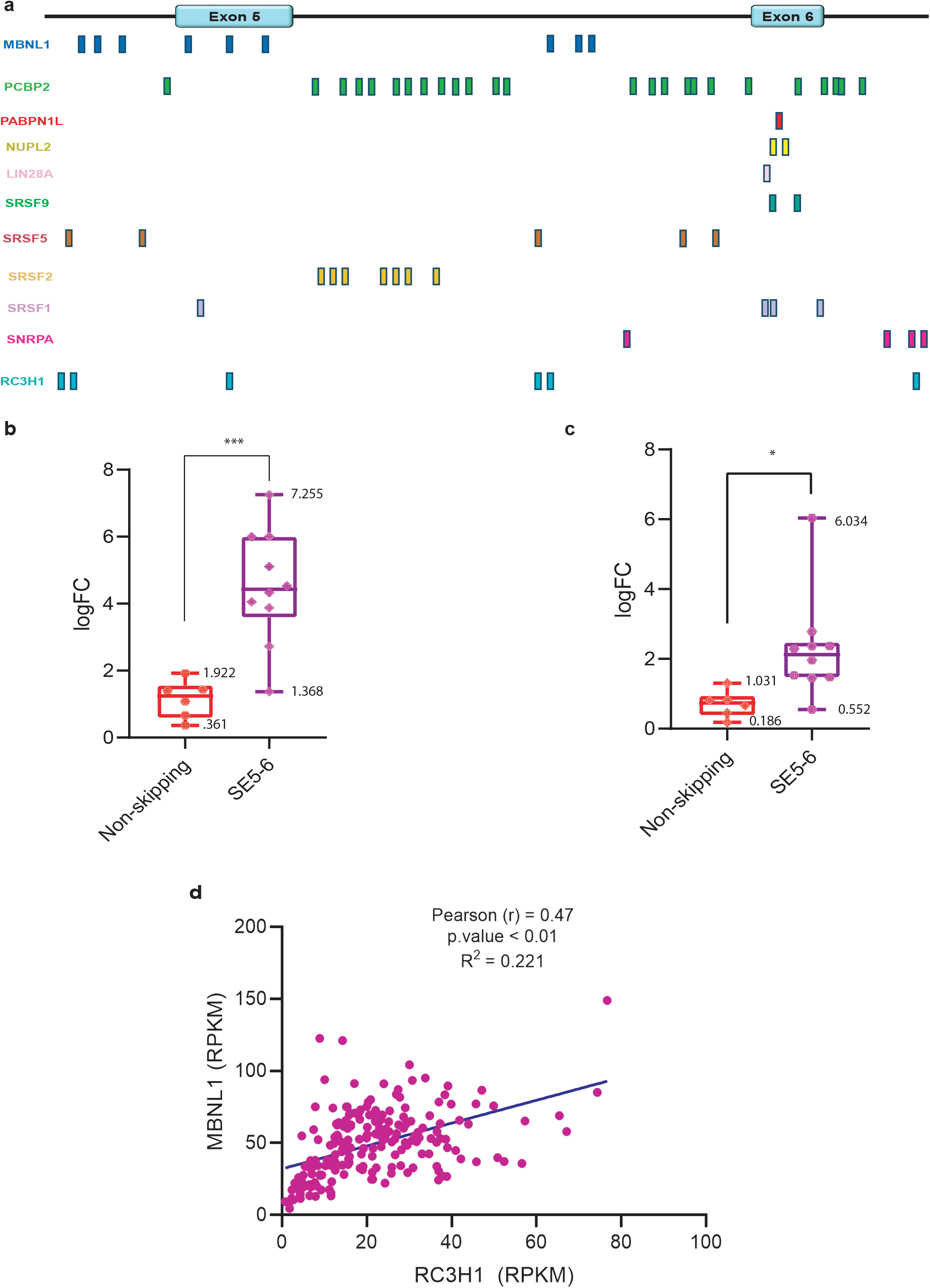
Identification of novel RBPs regulating CD19 exon 5-6 skipping. (a) Common RBPs obtained from public databases RBPmap and oRNAment along with their binding sites (from RBPmap-UCSC genome browser’s data) on CD19 exon 5-6 locus. (b) qRT-PCR showing that RC3H1 is significantly upregulated (unpaired t-test, p.value < 0.01) in B-ALL patient samples exhibiting skipped exon 5-6 (SE5-6) transcripts (n =10) when compared to samples showing no skipping bands (n = 6). (c) qRT-PCR showing that MBNL1 is significantly upregulated (unpaired t-test, p.value < 0.05) in B-ALL patient samples exhibiting skipped exon 5-6 (SE5-6) transcripts (n =10) when compared to samples showing no skipping bands (n = 6). Herein, S5, S13, S16, S18, S20, S25, S27, S36, S38, S44 were taken as SE5-6 samples whereas S19, S21, S22, S30, S37, S42 samples were taken as non-skipping samples. (d) Correlation analysis of RNA-Seq RPKM gene expression data extracted from “Pediatric Acute Lymphoid Leukemia - Phase II (TARGET, 2018)” deposited in cBioPortal, performed using GraphPad prism, shows that RC3H1 and MBNL1 are moderately correlated with each other (Pearson coefficient (r) = 0.47, p.value < 0.01).

## Discussion and conclusion

Alternative splicing plays a crucial role in regulating proteome diversity. Alterations in the splicing process results in the production of aberrantly spliced isoforms which are known to drive disease progression and drug resistance ^25,26^. Tumor cells exploit the presence of these irregularly spliced isoforms as a strategy to evade anti-cancer treatments ^27,28^. CD19 is a signaling antigen which plays a significant role in B-cell development, making it an attractive target for CART therapy ^29^. CART therapy has shown immense promise in treating B-ALL patients but a significant percentage of patients exhibit aggressive relapse of B-ALL post CART treatment ^30^. Aberrant splicing of CD19 which leads to a truncated CD19 protein has been identified as a factor contributing to resistance against CART therapy ^26^. The primary objective of our study was to conduct a comprehensive examination of CD19 splicing in primary patient samples and get insights into the presence of altered CD19 isoforms in patients prior to any therapy. Our results demonstrate that SE5-6 variant of CD19 is present in primary pediatric B-ALL patients irrespective of the subtype. Exon 5 and exon 6 engender transmembrane and cytosolic domains of CD19 respectively, therefore, an isoform that lacks these exons will not possess the normal CART epitope, enabling it to avoid detection by CARTs. In our study, we observed that skipping of exon 5-6 in CD19 results in generation of a pre-mature stop codon leading to absence of transmembrane & cytosolic domains in CD19. Additionally, it has been shown that phosphorylation of tyrosine residues (Y482 and Y513) within the cytosolic domains of CD19 regulates intracellular signaling cascades essential for B-cells maturation and survival, highlighting the importance of cytosolic domain for normal B-cell development ^31^.

Mechanistically, RBPs control alternative splicing program by interacting with cis-acting elements within the pre-mRNA, thereby regulating exon skipping and/or inclusion. Consequently, any alteration in RBP expression can lead to loss of target CD19 epitope, as suggested by earlier studies as well ^17,18,23^. Our in-silico analysis identified an extensive group of RBPs that were dysregulated in B-ALL patients and were predicted to have good affinity with CD19 exon 5-6 loci. Intriguingly, our complementary in-vitro analysis using qRT-PCR suggests that RC3H1 and MBNL1 expression levels is associated with exon skipping, as both the RBPs were significantly upregulated in patient samples exhibiting SE5-6 isoforms when compared to non-skipping samples. Furthermore, analysis of TARGET-phase II pediatric ALL dataset deposited on cBioPortal indicates that RC3H1 and MBNL1 are moderately correlated with each other, which aligns with our in-vitro findings. Conversely, both PCBP2 and SRSF2 exhibited non-significant differences in expression in qRT-PCR analysis and showed insignificant correlation with MBNL1 & RC3H1, respectively. Extensive research has highlighted the role of MBNL1 in B-ALL pathogenesis, as evidenced by numerous previous studies. MBNL1 knockdown promotes skipping of CD19 exon 2 ^23^ and diminishes levels of B-ALL markers genes has been observed previously ^32^. In addition, MBNL1 regulates aberrant splicing of genes involved in KMT2A rearranged oncogenesis has also been established ^33^. Interestingly, a recent retrospective study reported that patients with KMT2A subtype have very poor overall survival post CART therapy relapse ^34^, this suggests that MBNL1 dysregulation may be associated with poor response towards CART therapy, directly or indirectly. Furthermore, RC3H1 has been found to hinder anti-tumor immunity, and mutations in RC3H1 have been linked to immune system dysregulation ^35,36^. In summary, our in-silico and in-vitro data indicates that RC3H1 and MBNL1 could be key biomarkers for CART resistance, highlighting the need for in-depth studies to confirm their role in exon 5-6 skipping.

Hence, to conclude, our study investigates the presence of aberrantly spliced CD19 isoform responsible for CART resistance, in a large cohort of primary pediatric B-ALL patients, and sheds light on the RBPs regulating this aberrant splicing. Patients having this isoform at the time of diagnosis are more prone to develop CART resistance as this isoform might become more dominant during therapy. In future, the presence of these isoform in the diagnosis stage itself, may be used as a screening criterion to provide more personalized or combination therapy to the patients.

## Acknowledgements

This work is supported by ICMR (2021-13073) and intramural grants from IIT Hyderabad to A.M. D.S. is a recipient of PMRF, GoI.

