## Supplementary for "Widespread Prevalence of CD19 Exon 5-6 Skipping In Indian Pediatric B-cell Acute Lymphoblastic Leukemia Patients"

Supplementary figure 1:

a

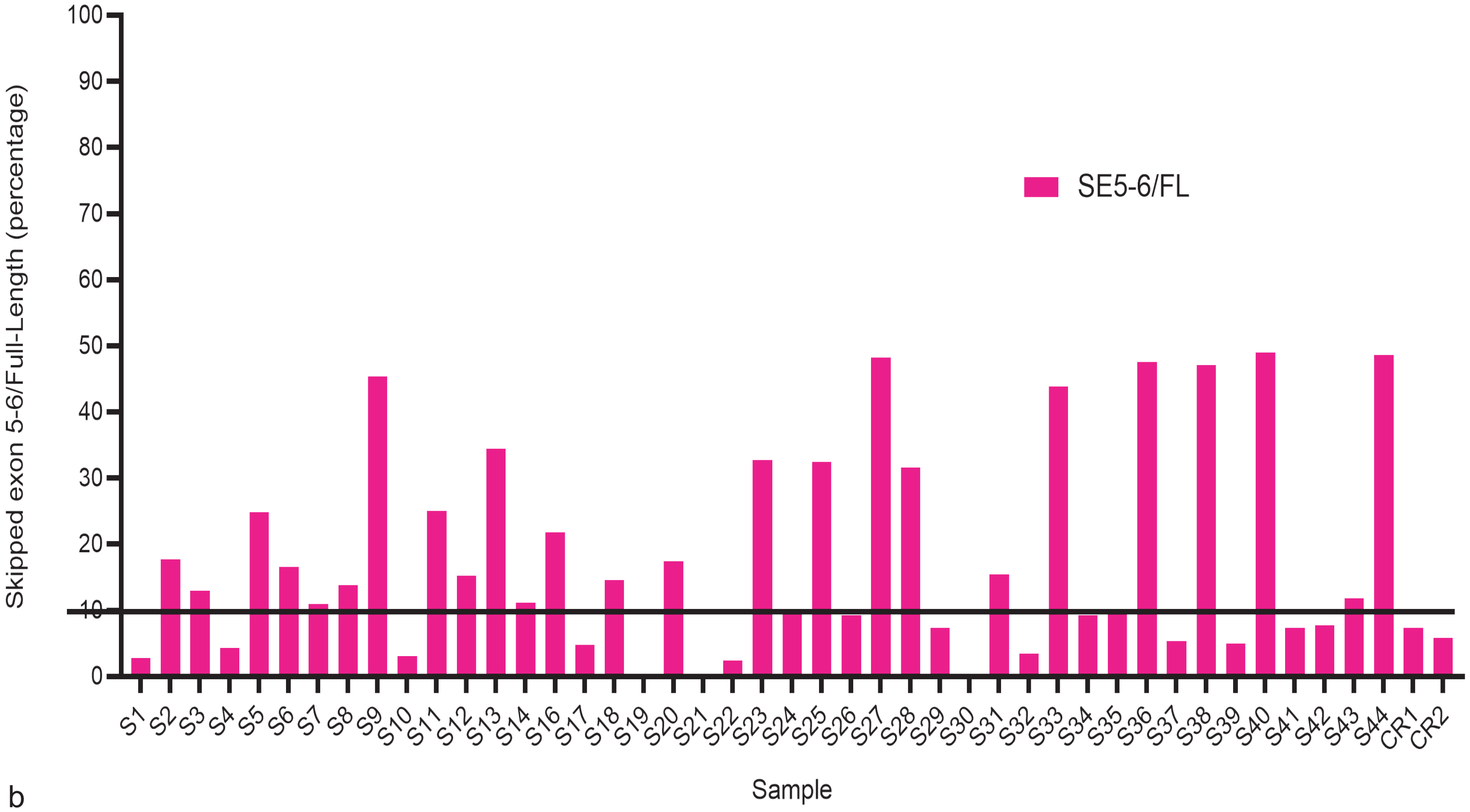


b

```
CD19FL          ---------------agmgatcgcccggcmgagatatgtgggtaatggagacgggtctgt   45
CD19exon4-8     GCCTAGAGCTGAAGGACGATCGCCCGGCCAGAGATATGTGGGTAATGGAGACGGGTCTGT   180
CD19SE5-6       -------------tagmgatcgcccggccagagatatgtgggtaatggagacgggtctgt   47
                                       **  *  * ******************************

CD19FL          tgttgccccgggccacagctcaagacgctggaaagtattattgtcaccgtggcaacctga   105
CD19exon4-8     TGTTGCCCCGGGCCACAGCTCAAGACGCTGGAAAGTATTATTGTCACCGTGGCAACCTGA   240
CD19SE5-6       tgttgccccgggccacagctcaagacgctggaaagtattattgtcaccgtggcaacctga   107
                ************************************************************

CD19FL          ccatgtcattccacctggagatcactgctcggccagwactatggcactggctgctgagga   165
CD19exon4-8     CCATGTCATTCCACCTGGAGATCACTGCTCGGCCAGTACTATGGCACTGGCTGCTGAGGA   300
CD19SE5-6       ccatgtcattccacctggagatcactgctcggcc--------------------------   141
                **********************************

CD19FL          ctggtggctggaaggtctcarctgtgactttggcttatctgatcttctgcctgtgttccc   225
CD19exon4-8     CTGGTGGCTGGAAGGTCTCAGCTGTGACTTTGGCTTATCTGATCTTCTGCCTGTGTTCCC   360
CD19SE5-6       ------------------------------------------------------------   141

CD19FL          ttgtgggcattcttcatcttcaaagagccctggkcctgaggaggaamakaaagcgaatga   285
CD19exon4-8     TTGTGGGCATTCTTCATCTTCAAAGAGCCCTGGTCCTGAGGAGGAAAAGAAAGCGAATGA   420
CD19SE5-6       ------------------------------------------------------------   141

CD19FL          ctgaccccaccaggagattcttcaaagtgacgcctcccccaggaagcgggccccagaacc   345
CD19exon4-8     CTGACCCCACCAGGAGATTCTTCAAAGTGACGCCTCCCCCAGGAAGCGGGCCCCAGAACC   480
CD19SE5-6       --------------agattcttcaaagtgacgcctcccccaggaagcgggccccagaacc   187
                              **********************************************
```

Supplementary figure 2:

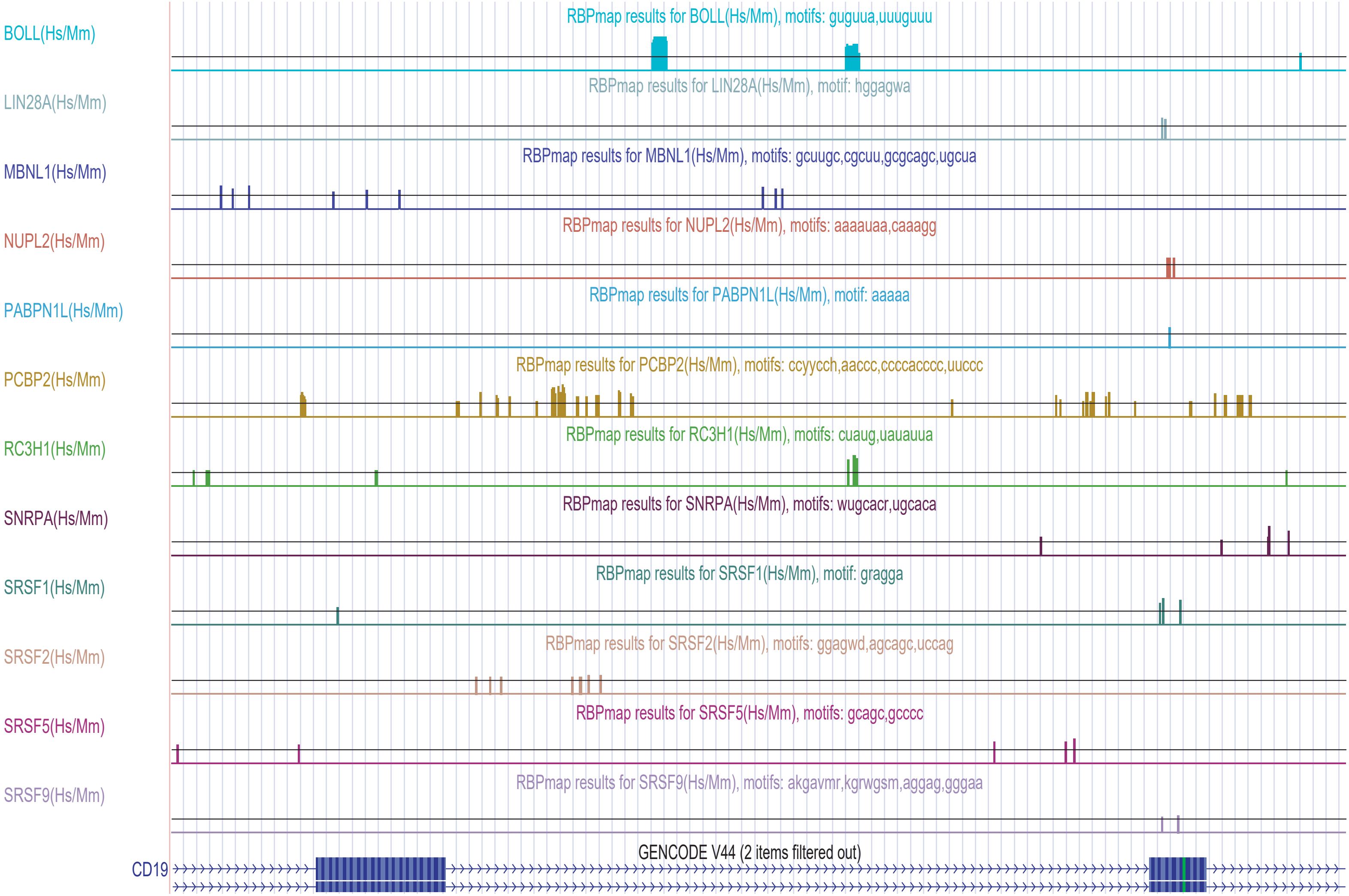

Supplementary figure 3:

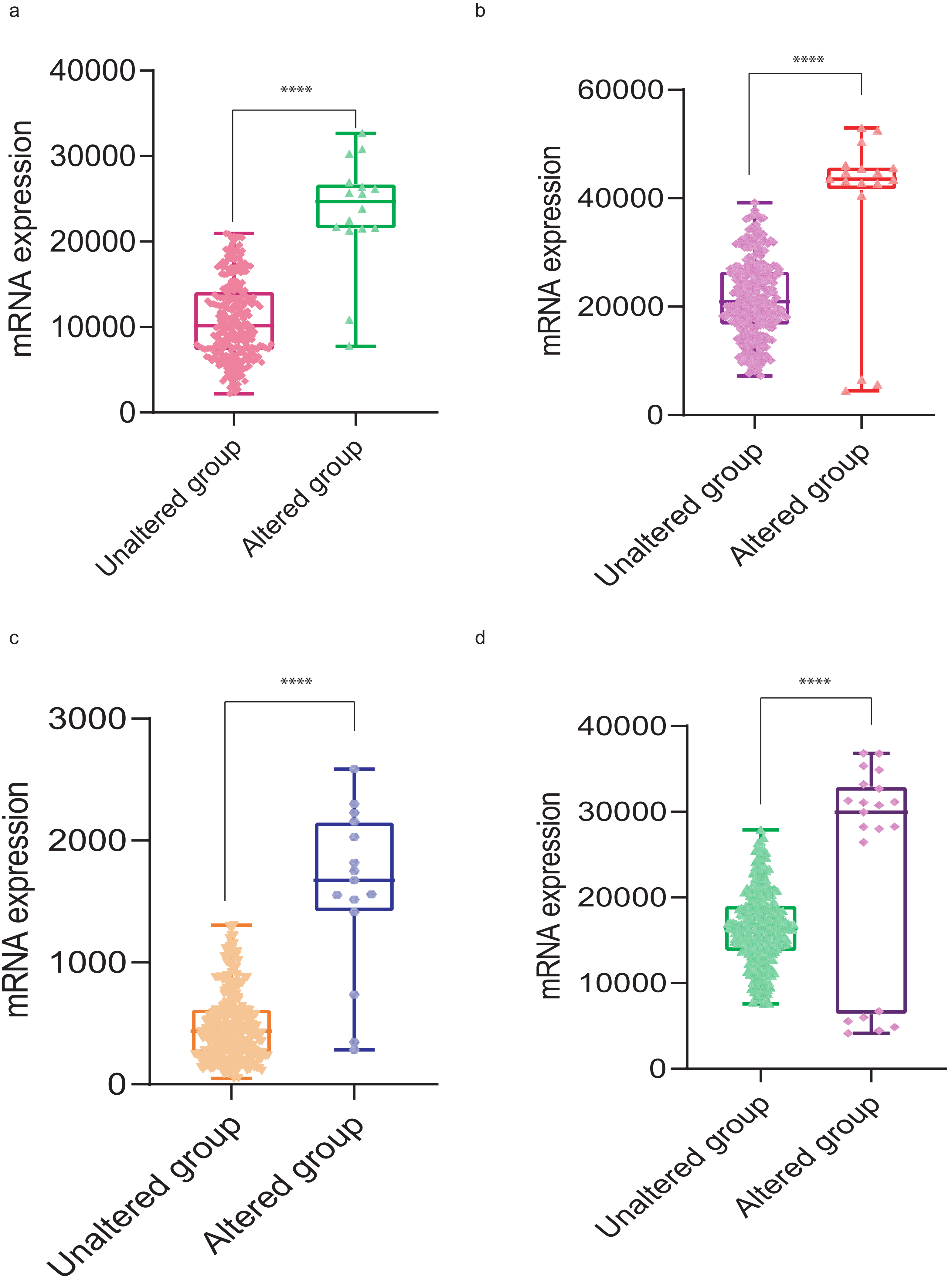

Supplementary figure 4:a

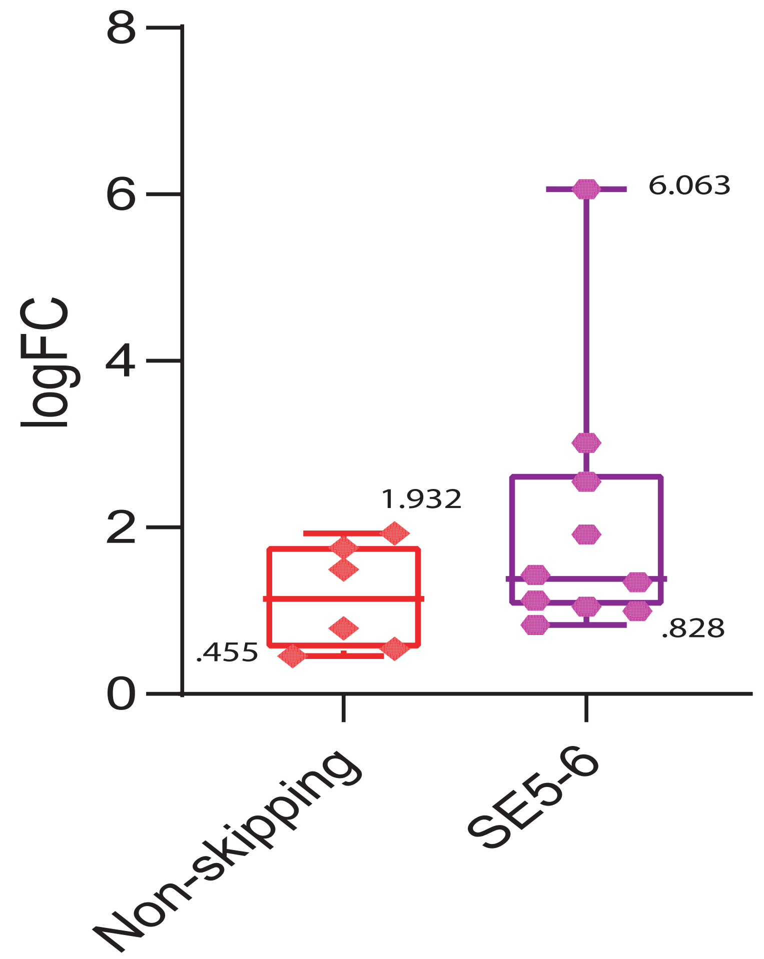


b

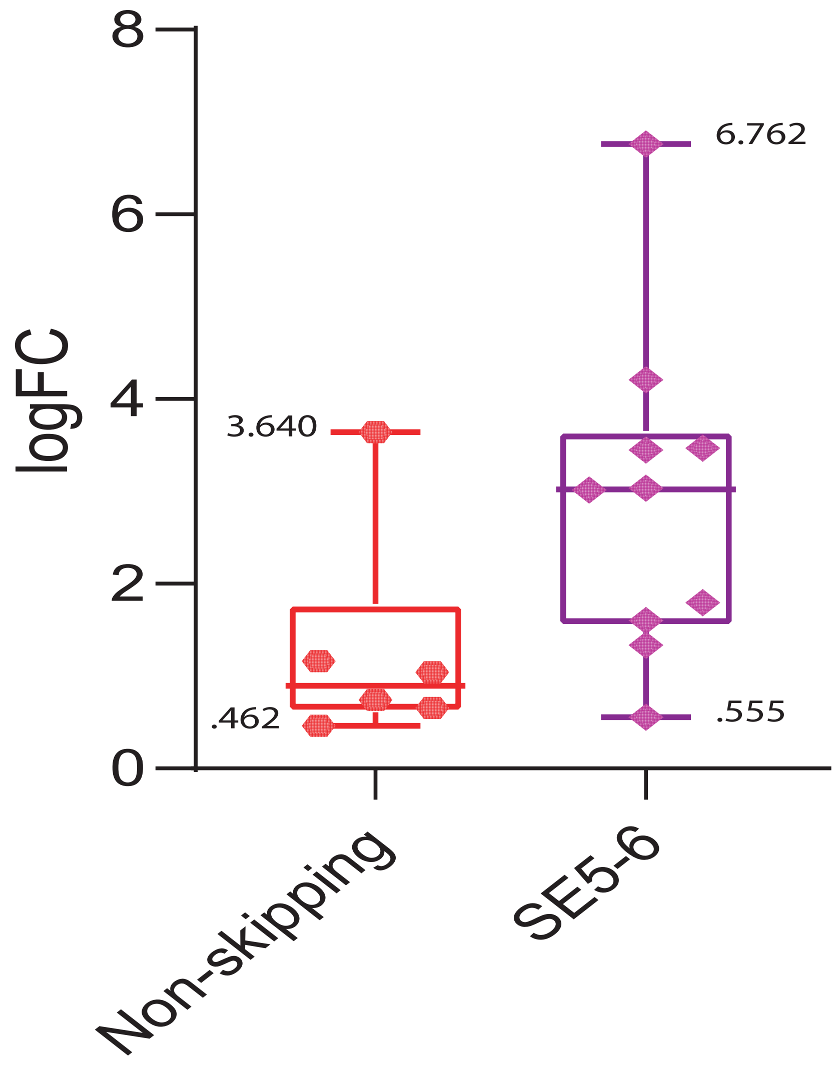
